# Micro-scale ecology regulates particulate organic matter turnover in model marine microbial communities

**DOI:** 10.1101/241620

**Authors:** Tim N. Enke, Gabriel E. Leventhal, Matthew Metzger, José T. Saavedra, Otto X. Cordero

## Abstract

The degradation of particulate organic matter in the ocean is a central process in the global carbon cycle, the ‘mode and tempo’ of which is determined by the bacterial communities that assemble on particle surfaces. Although recent studies have shed light on the dynamics of community assembly on particles –which serve as hotspots of microbial activity in the ocean, the mapping from community composition to function, i.e. particle degradation, remains completely unexplored. Using a collection of marine bacteria cultured from different stages of succession on chitin micro-particles we found that the hydrolytic power of communities is highly dependent on community composition. Different particle degrading taxa –all of which were early successional species during colonization– displayed characteristic particle half-lives that differed by ~170 hours, comparable to the residence time of particles in the ocean’s mixed layer^1^. These half-lives were in general longer in multispecies communities, where the growth of obligate cross-feeders limited the ability of degraders to colonize and consume particles. Remarkably, above a certain critical initial ratio of cross-feeder to degrader cells, particle degradation was completely blocked along with the growth of all members of the community. We showed that this interaction occurred between a variety of strains of different taxonomic origins and that it only appears when bacteria interact with particles, suggesting a mechanism by which non-degrading secondary consumers occlude access to the particle resource. Overall, our results show that micro-scale community ecology on particle surfaces can have significant impact on carbon turnover in the ocean.

## Introduction

Learning how the composition of ecological community impacts their function is arguably one of the central challenges in ecology^2–4^. In the case of microbes, this problem is particularly complex, not only because of the extreme diversity of taxa and genes that make up microbial communities, but also because community function depends on micro-scale processes that are hard to measure such as aggregation, dispersal and cell-cell interactions^5^. A prime example of the link between micro-scale community ecology and large-scale ecosystem function is found in the biological turnover of particulate organic matter. In the marine environment, biopolymer particles formed by aggregation of fragments of decaying organisms, fecal pellets, and extracellular polysaccharides are degraded and consumed by heterotrophic bacteria that attach to particle surfaces and form dense microbial communities of large taxonomic and metabolic diversity^6–9^. Because particulate matter tends to sink in the water column, its degradation in the upper layers of the ocean where oxygen abounds is crucial to sustain the marine food web and prevent the sequestration of carbon and nitrogen into the deep sea^9–11^. Therefore, particle-attached microbial communities play a fundamental role by closing the loop of the global carbon cycle and maintaining the balance of nutrients in marine ecosystems. Although many physical aspects of the bacteria-particle interaction such as attachment or the effects of flow^12,13^ have been well characterized, the possible role that ecological interactions between microbes may play in controlling the dynamics of particle colonization and degradation –and thus the ‘mode and tempo’ of the global carbon cycle– is much less clear.

Previous studies have shown that ecological interactions between microbes can play a significant role in controlling the dynamics of community assembly on particles. Competition for particle surface and thus primary resource access is likely to be strong among particle-attached bacteria and interference competition mediated by secondary metabolites can be a powerful strategy to deter competitors^14,15^. Moreover, over the time scales of particle turnover, trophic interactions mediated by byproducts of degradation and primary metabolism can strongly influence the overall dynamics of bacterial growth^16^: To release the carbon trapped in particulate matter, bacteria secrete hydrolytic enzymes that deconstruct complex biopolymers and release soluble sugars into the environment. The bioavailable sugars can in turn be taken up by nearby cells, thus unlocking a niche for ‘cheaters’ that consume resources but do not contribute to degradation^16,17^. Likewise, byproducts of primary metabolism such as organic acids or amino acids that are released to the local environment can be consumed by crossfeeding bacteria that co-assemble on the particle. On chitin particles, these types of trophic interaction have been shown to lead to successional waves and invasion of secondary consumers, which eventually become the numerically dominant members of the community^16^. These findings led us to hypothesize that interactions across trophic levels at the micro-scale might alter the catabolism of chitin and consumption of byproducts, possibly affecting the rate of particle turnover and the conversion from particle to bacterial biomass.

To test this hypothesis, in this study we used an isolate collection obtained directly from particle-attached communities previously shown to colonize in micro-scale successions^16^. In brief, these communities were enriched on ~50 μm paramagnetic chitin hydrogel particles incubated in seawater from the coastal ocean (Nahant, MA, USA). Bacteria were isolated directly from the particles, resulting in a collection that includes taxa such as *Alteromonadales, Flavobacteriaceae, Rhodobacteriales, Vibrionaceae*, and *Oceanospiriliae*. Notably, the composition of our collection coincides well with the taxonomic profiles of natural chitinous marine particles collected at 200–500 meters depth in the North Pacific gyre^18^. This overlap between our isolate collection and the taxonomic composition of natural particle-attached communities suggests that isolates obtained from model particles represent a relevant set of strains with which to study the effect of ecological interactions on particle turnover.

Bacterial isolates in our collection fall into two coarse-grained functional groups, defined on the basis of shared physiological characteristics and colonization dynamics^16^. The first group comprises *primary degraders*, which secrete chitinolytic enzymes, are motile, can grow rapidly on degradation byproducts and belong to species that tend to appear early during particle colonization. The second group corresponds to *secondary consumers*, which in general do not secrete enzymes, cannot grow on chitin, grow poorly if at all on monomers, are not motile and tend to belong to late successional species (Fig 1A, Fig S1). Although secondary consumers cannot grow on chitin particles alone, they can reach 100–1000 fold higher abundance in the presence of primary degraders^16^ due to their ability to utilize metabolic byproducts released by primary degraders during colonization.

**Figure 1:**
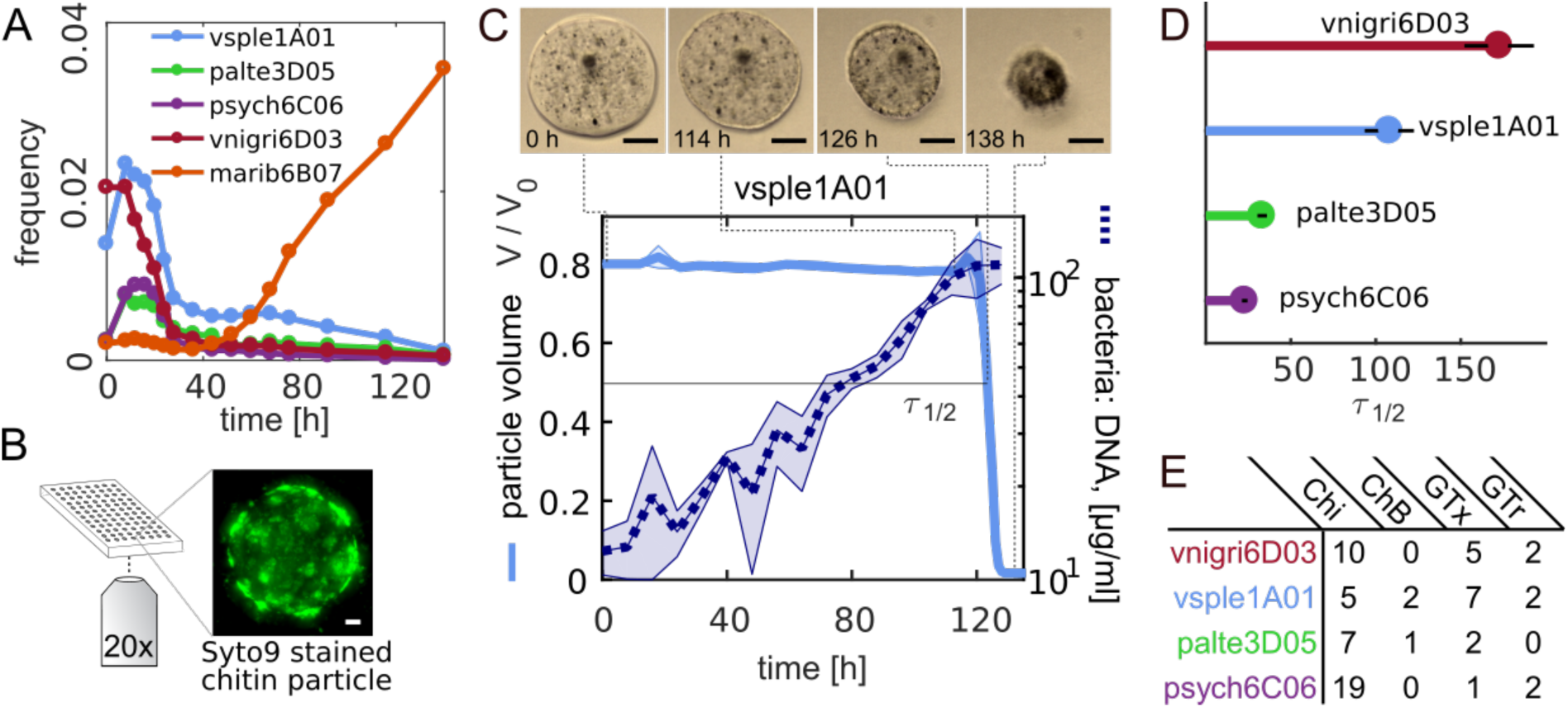
Particle degradation dynamics of bacteria isolated from chitin micro-particles. **A)** Culture independent dynamics of four primary degraders (vsple1A01, palte3D05, psych6C06, vnigri6D03) and a secondary consumer (marib6B07). Trajectories shown depict dynamics of selected taxa in particle incubations with raw seawater. Data from ref. 16. **B)** In the laboratory, chitin particles immersed in bacterial suspensions are imaged at the bottom of microtiter plates for up to 240 h. The particle image corresponds to DNA stained palte3D05 after 24 h, showing the formation of bacteria micro-colonies on the particle surface. Scale bar corresponds to 10 μm. For a 3D animation of the image, see supplemental video 1. **C)** Upper panel: Phase contrast (20x) micrographs of a chitin particle cross section taken at different time points during incubation with vsple1A01. Scale bar: 30 μm. (See also supplementary video 2). Lower panel: Particle volume over time normalized to initial volume (solid line) and bacterial abundance as measured by the amount of DNA extracted from ~100 particles at different points of colonization (dashed line). The standard deviation of measurements was calculated using three replicate particles from the same well, and three different bulk incubations for DNA. **D)** Particle half-lives for the four different degraders tested with an inoculum of ~5×10^5^ cells per ml. **E)** Number of gene copies of chitinases (Chi), chitin binding proteins (ChB), GlcNAc specific chemotaxis (GTx) and transport (GTr) genes. See also Supplementary Table 1.

Our goal in this study is to provide a quantitative description how particle degradation kinetics depend on the assembly of primary degraders and secondary consumers during particle colonization. To this end, we first studied how mono-cultures of primary degraders consumed particles by tracking changes in particle volume over time using high-throughput, high-resolution time-lapse microscopy (Fig 1B) and guiding our analysis with simple mathematical models of colonization and resource consumption. Subsequently, we assembled two-strain communities of primary degraders and secondary consumers and developed a quantitative phenomenological characterization of the impact of secondary consumers on degradation. Our results reveal that early colonizing taxa can differ significantly in their hydrolytic power to break down chitin, that particle degradation is limited by the number of enzyme-secreting bacteria that colonize the particle surface, and that secondary consumers effectively become parasites that increase in abundance at the cost of the primary degraders when co-colonizing on particle surfaces. Furthermore, the presence of parasitic secondary consumers can delay or even obstruct particle degradation. All these effects suggest that micro-scale community ecology on particle surfaces plays a major role in controlling community function by primarily slowing down resource turnover rates.

## Results

### Variability in hydrolytic power: the effect of primary degrader identity and abundance

We tracked the dynamics of particle consumption by measuring changes in particle volume over time, *V* (*t*), using high-throughput time-lapse microscopy of individual chitin microbeads. We chose an initial concentration of degrader cells of 5×10^5^ cells/ml –an upper-bound estimate of the concentration of degrading bacteria in coastal waters^19^- and quantified *V* (*t*) over a period of 240 h, for four primary degraders and four secondary consumers incubated in media with no carbon source other than the particle. As expected, secondary consumers did not grow on particles in monoculture and therefore did not affect *V*(*t*) over the course of the ten-day time-lapse. For primary degraders, instead, *V*(*t*) was characterized by a long period of no detectable change, followed by a swelling of the particle and an abrupt collapse (Fig 1C, Sup. movie 2). Measurements of bacterial growth during degradation showed that bacteria grew steadily from the beginning of the incubation, despite no apparent change in particle volume, indicating that depolymerization was a continuous process and that swelling and collapse occurred only after a critical amount of polymer was consumed (Fig 1C). Particle swelling indicates that the degradation of cross-linked chitin in the hydrogel allows water molecules to expand the matrix^20^, while the transition from swelling to collapse indicates the point at which depolymerization ‘outcompetes’ swelling. The type of degradation curves observed for primary degraders (Fig. S2), with most of the dynamics concentrated on long transients, allowed us to quantify the ability of bacteria to consume particles with a single quantity, the particle half-life, *τ*_1/2_, i.e. the time it took for the particle to decrease to half its volume (see methods).

We found a remarkable variation in *τ*_1/2_ among the four different primary degraders, despite the fact that all of these isolates appeared early on in the ecological succession on chitin particles (Fig 1A). At an initial cell concentration of 5×10^5^ cells/ml for all primary degraders, particle half-lives varied from ~30 h for the fastest degrader (a strain of the genus *Psychromonas*, named psych6C06) to ~200 h for the slow degraders (a strain of *Vibrio nigripulchritudo* named vnigri6D03) (Fig 1D). The large number of chitinase copies in psych6C06 (19 copies) suggested that gene dosage played a role in controlling the hydrolytic power of the strains. However, overall the differences between *τ*_1/2_ among primary degraders could not be clearly correlated to variation in gene content, suggesting instead that expression levels and the ‘quality’ of extracellular enzymes played a more significant role. Gene content did however distinguished primary and secondary consumers: degraders tended to encode the genomic potential to transport chitin monomers (N-acetylglucosamine specific PTS transporters), use monomers as chemotaxis signals and attach to chitin surfaces, features which tended to be absent in secondary consumers (Fig 1E, Table S1).

Chitin degradation is intrinsically linked to the production of public goods such as chitinases and as such can be subject to cooperative growth dynamics^21^, i.e. a positive dependency between cell densities and growth or depolymerization rates. If cooperativity does play a role, half degradation times would be highly sensitive to cell numbers, increasing disproportionally in cases where cell load is low. To test the relevance of this phenomenon and, in general, to study how *τ*_1/2_ depended on initial conditions, we measured degradation kinetics as a function of the initial concentration of primary degrader [*B_p_*]_0_, which until now was arbitrarily set to 5×10^5^ cells/ml. In addition, we guided our analysis with simple models of particle degradation and bacterial growth. To construct these models we assumed that particle depolymerization was proportional to the density of bacteria. We studied two possibilities, i) that bacteria grew cooperatively, i.e. with growth rate proportional to *B^n^* and *n* > 1, and ii) that cooperativity played no significant role and growth and occurred at fixed, density independent per capita rates. Assuming that *τ*_1/2_ depends linearly on the speed of depolymerization, model i) predicts that *τ*_1/2_ should scale as − **1**/[*B_p_*]_0_, whereas model ii) predicts that *τ*_1/2_ should scale as −log ([*B_p_*]_0_) (Methods and Supplementary Text, Fig. 2F).

In agreement with the simplest model with no cooperativity (ii), we find a linear relation between *τ*_1/2_ and log ([*B_p_*]_0_) (Fig 2, Table S2). This behavior implies that the particle half-life is controlled by simple mass action kinetics^22^ that—at least in the conditions of our experiment—are not influenced by cooperativity. More precisely, we find that *τ*_1/2_ is well described by the following expression,
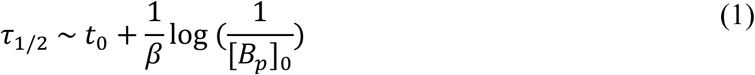

where *t_0_* is the intercept of the lines in Fig 2E and represents a timescale to degradation that is intrinsic to each strain, *β* is the slope and represents the per-capita contribution to the degradation process and –log 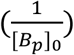 captures the effect of the primary degrader concentration in the local environment, akin to a chemical potential for the cell-particle reaction.

**Figure 2.**
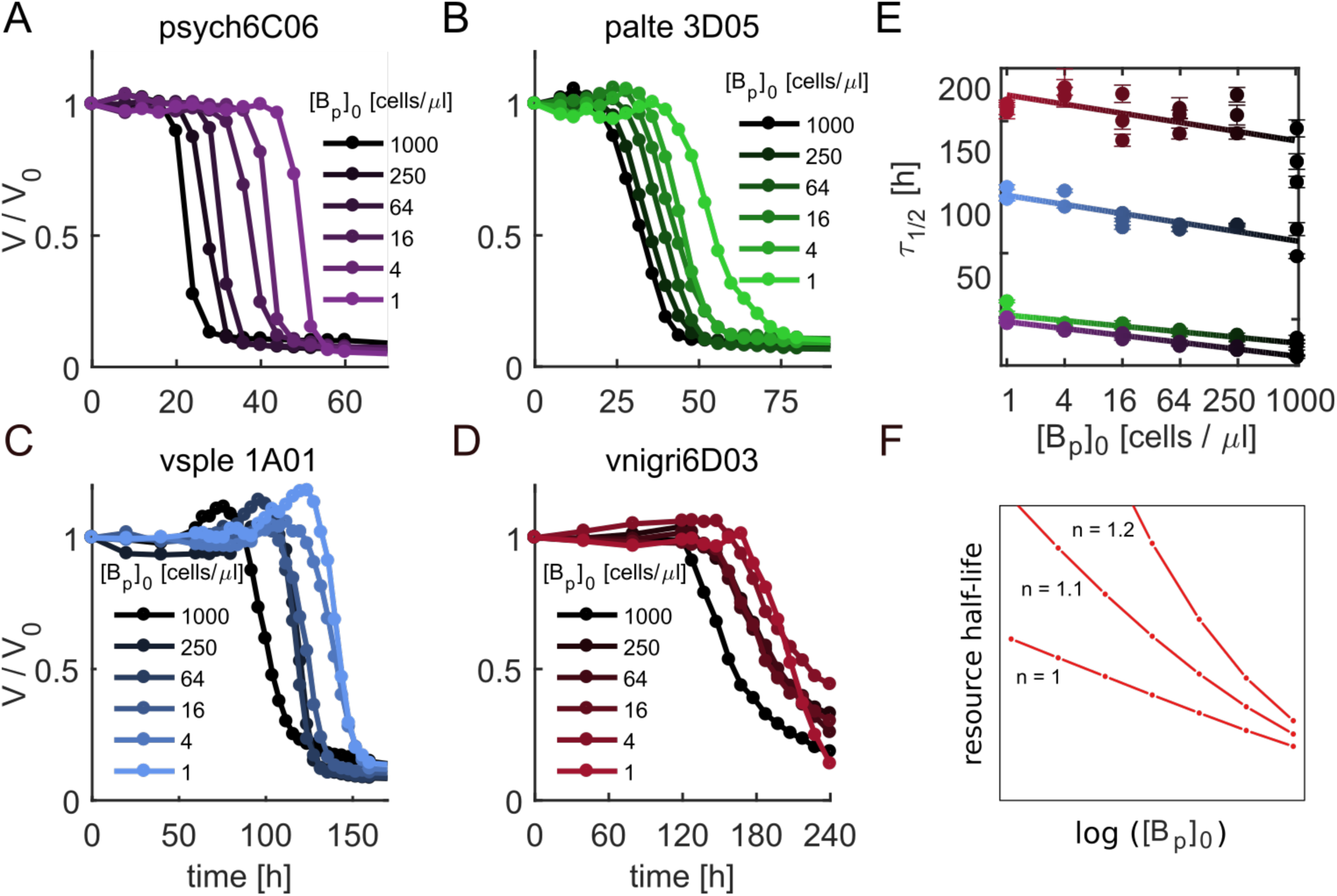
Effect of cell initial cell concentration on particle degradation kinetics. **A-D)** Mean particle volume over time for primary degraders, over a range of initial inoculum concentrations, [B_p_]_0_ (See Fig S3). **E)** Linear dependency between the log_2_([B_p_]_0_) and the particle half-life as predicted by equation (1) validating the simple model of degradation without cooperativity (See also Table S2). **F)** Prediction for log([B]_0_) vs half-life based on models with (n > 1) or without cooperativity (n = 1).

The relationship found in (1) shows that the turnover of particulate organic matter depends on the load of primary degraders in the *milieu* in a simple, predictable manner. The lack of a cooperativity observed suggests that the possible benefits that bacteria may derive from ‘teaming up’ are effectively offset by local competition for resources between neighbors. Overall, our results indicate that variation in the composition and abundance of primary degraders can have a significant impact on the rate of particulate organic matter turnover.

### Secondary consumers behave as parasites during particle degradation

To understand how ecological interactions between primary degraders and secondary consumers influence particle degradation, we focused our analysis on two primary degraders and one secondary consumer. We chose the relatively ‘slow’ degrader, vsple1A01 (Fig 1CD) a member of the *Vibrio splendidus* clade, the most abundant group of marine vibrios in coastal seawaters^23^, and the relatively ‘fast’ degrader, *Pseudoalteromonas sp*. palte3D05 (Fig S2), a common member of heterotrophic bacterioplankton communities^24,25^. Secondary consumers, or strains unable to degrade chitin, have previously been found to invade particle-attached communities and to become numerically dominant during community assembly^16^ (Fig. 1A). We focused our efforts on a secondary consumer cultivated from seawater-incubated chitin particles, a strain of the genus *Maribacter* (a type of marine *Flavobacteria*), that we here call marib6B07. As with other secondary consumers, marib6B07 is able to crossfeed when grown in co-culture with degraders^16^. Interestingly, genome sequences marib6B07 and other secondary consumers show that, despite their inability to degrade chitin under laboratory conditions, these organisms can contain chitinases (marib6B07 has two), but in general lack genes for *N*-acetylglucosamine specific chemotaxis, *N*-acetylglucosamine specific phosphotransferase (PTS) transport and chitin-binding, all of which tend to be present in multiple copies in the genomes of primary degraders (Table S1). These differences in the genomes of primary degraders and secondary consumers suggest that their functional roles in the community may be determined by the interaction between multiple traits, such as the ability to chemotax towards breakdown products of chitin and to transport them into the periplasm.

Co-incubation of mari6B07 with vsple1A01 and palte3D05 showed that mari6B07 increased *τ*_1/2_ relative to primary degrader monocultures (Fig 3A), implying that the crossfeeder impaired the ability of degrader populations to depolymerize the particle. To study this phenomenon in a quantitative manner, we measured how *τ*_1/2_ responded to changes in the initial concentration of secondary consumer, [*B_s_*]_0_, with the number of cells of the primary degrader fixed at a given concentration ([*B_p_*]_0_ ≈ 1.25×10^5^ cells/ml) (Fig 3A, Fig S5A). We found that over low [*B_s_*]_0_, *τ*_1/2_ increased roughly linearly, such that a one-fold increase in the secondary consumer [*B_s_*]_0_ had approximately the same effect as a ten-fold reduction of the primary degrader [*B_p_*]_0_ in monoculture.

**Figure 3:**
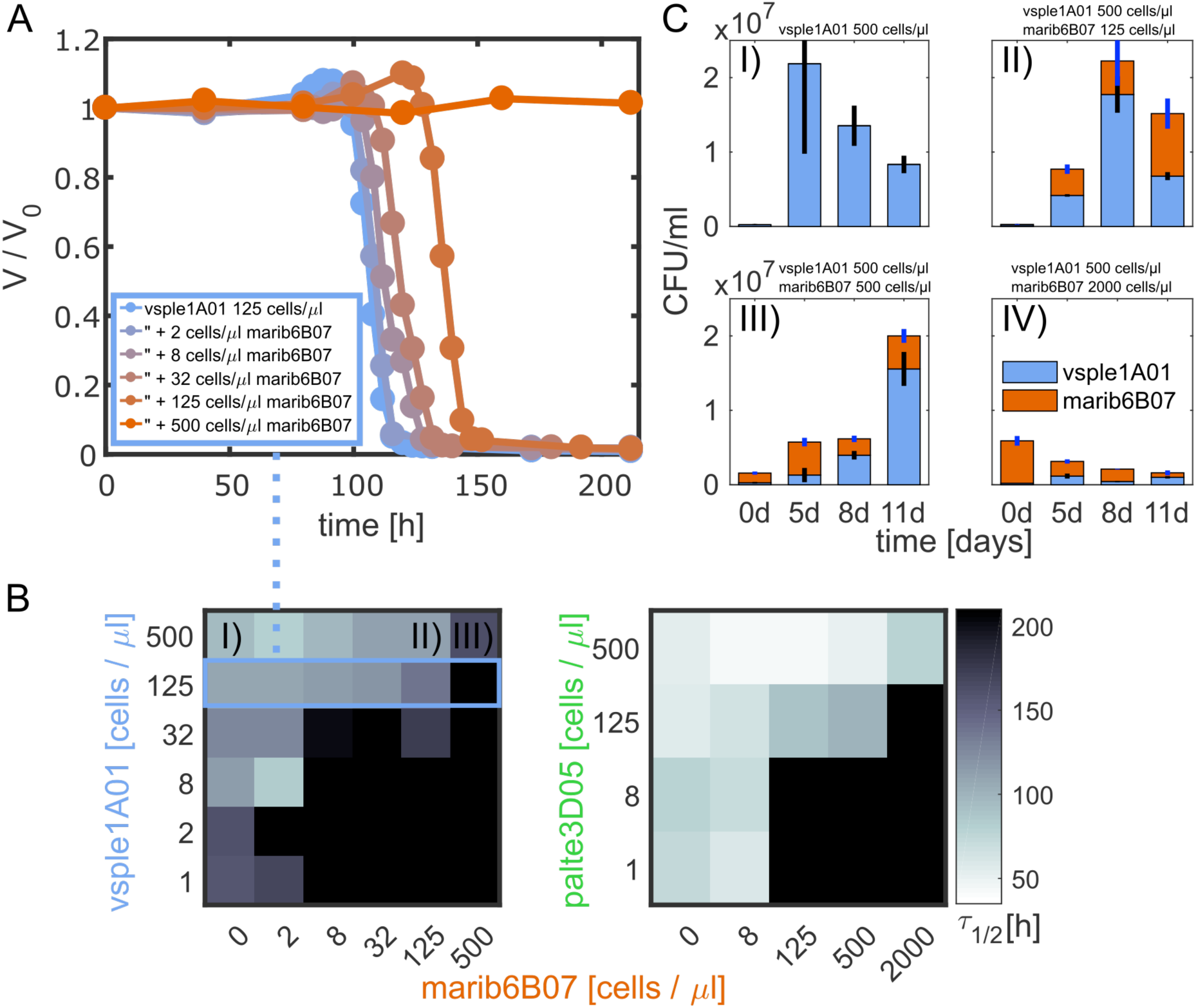
Secondary consumers inhibit degradation. **A)** Particle degradation curves with different marib6b07 concentrations. At increasing concentrations of the secondary consumer the particle halflife increases disproportionally beyond the 220 h time limit. **B)** Heat maps depict *τ*_1/2_ as a function of different primary degrader and secondary consumer inoculum concentrations and show that particle half-lives depend on the relative concentrations of primary degrader and secondary consumer cells. Color scale is the same for both heat maps. The blue highlighted row of the heat map corresponds to the degradation curves in A). For all degradation curves used for the heat maps see FigS6 and FigS7, respectively. **C)** CFUs of vsple1A01 and marib6B07 during co-culture on chitin particles, showing that marib6B07 acts as a parasite that grows “at the expense” of vsple1A01’s yield. I) vsple1A01 in mono-culture. Particle degradation observed at ~5 days. II,III) co-cultures, particle degradation observed at ~8 d and ~11 d, respectively. IV) co-culture: no degradation observed (standard deviations for N=3 replicates). Decrease in CFUs is due to loss of viability after degradation.

Surprisingly, at a threshold [*B_s_*]_0_ we observed an abrupt increase in *τ*_1/2_, to the extent that particle degradation did not occur within the 240 h imaging period, suggesting that the population of primary producers might have been inhibited from colonization and/or growth. To investigate how this phenomenon depended on the composition of the two-strain community, we varied the abundance of the primary degraders, [*B_p_*]_0_ and secondary consumer [*B_s_*]_0_, in order to obtain degradation phase planes (Fig 3B). The degradation phase planes show that complete inhibition did not depend on the total concentration of the secondary consumer, [*B_s_*]*_0_*, but on the ratio of secondary consumer to primary degrader, *γ* = [*B_s_*]_0_*/*[*B_p_*]_0_ (Fig 3B,C). For the slow degrader, vsple1A01, degradation was blocked at *γ* > ~ 1, whereas for the fast degrader, palte3D05, degradation was blocked above a ratio of *γ* > ~16, showing that the slow degrader was more sensitive to the inhibitory effects of secondary consumer marib6B07 than the fast degrader. This analysis indicates that the balance between the relative abundances of secondary consumers to primary degraders in the environment, in addition to the degradation kinetics of the primary consumer, may be an important parameter that dictates the turnover rates of carbon over short time-scales (see Discussion).

Quantification of the abundance of each strain in co-culture before and after particle degradation showed that the interaction between primary degrader and secondary consumer is parasitic i.e. positive for the consumer, negative for the degrader. CFU counts during the time course of degradation in co-cultures of vsple1A01 and marib6B07 showed that primary degrader growth rate and yield were lower than in monoculture, and that the “loss” of degrader cells was compensated by the growth of secondary consumers (Fig 3C). Secondary consumers doubled approximately 5 times by the time of particle collapse, in contrast to their zero doublings in monoculture (see Fig S5B). Notably, the total yield of the co-culture was always equal or lower to the yield of the mono-culture, highlighting the parasitic nature of the interaction. Thus, secondary consumers, whose growth is facilitated by primary degraders, exert a negative feedback on degraders, limiting their ability to consumer produced resources and potentially their own growth.

Given the higher ratio of secondary consumer to degrader (*γ*) required to inhibit palte3D05 compared to vsple1A01, we hypothesized that “slow degraders” might be more susceptible to the detrimental effect of secondary consumers. To test this hypothesis as well as whether the observed parasitic interactions can be generalized to other primary degrader – secondary consumer pairs, we measured the effect of co-culture at *γ* = 1 ratio on particle degradation for all primary degraders (Fig 1D) with four different secondary consumers (including marib6B07) of diverse taxonomic origins, all of which were co-isolated from the same chitin-attached communities. The results showed that while the fast degraders psych6C06 and palte3D05 were only mildly affected by co-culture with secondary consumers at *γ =* 1, the slow degraders vsple1A01 and vnigr6D03 were susceptible to the presence of secondary consumers (Fig 4), with the slowest degrader, vnigr6D03 being inhibited by all four secondary consumers, three of which caused total blockage of particle consumption. These data further indicate that parasitic interactions between degraders and consumers are not dependent on specific taxa, but rather on the hydrolytic power of the degrader.

**Figure 4:**
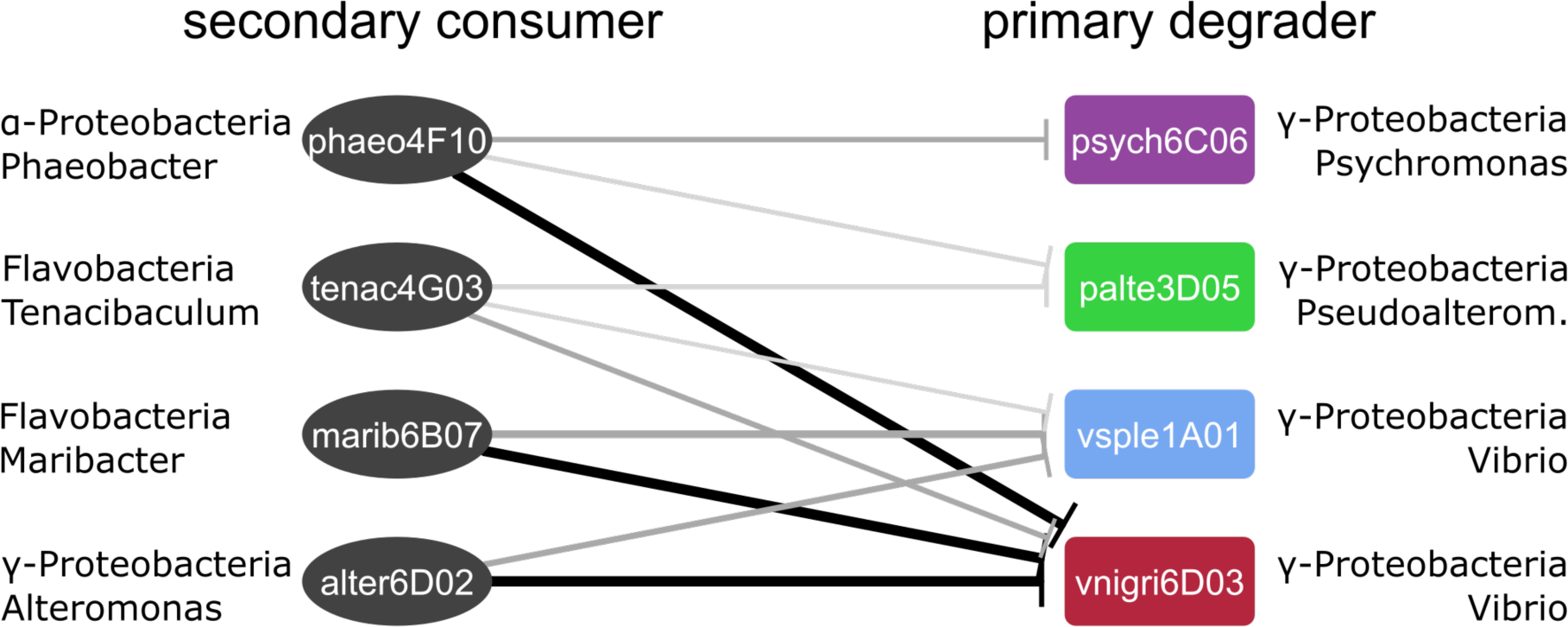
Degradation inhibition is specific to functional groups, not strains. The network depicts the effect of four different secondary consumers from diverse taxonomic origins on the characteristic half live *τ*_1/2_ during particle degradation by primary degraders with different hydrolytic power). Network edge width is proportional to 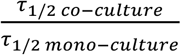. Edges are drawn between secondary consumers and primary degraders when the mean (n=3) half lives *τ*_1/2_ determined from the degradation curves of mono- and co-cultures were determined to be statistically different by one-way ANOVA. Black: complete inhibition, no *τ*_1/2_ determined in co-culture, dark grey: p < 0.05, light grey: p < 0.1, respectively. See Figure S9 for raw data.

Consistent with the observation that interactions are not specific to strains or species but to functional roles (i.e. secondary consumer, primary degrader), we did not find evidence of chemical antagonism from secondary consumers to degraders. Agar plate assays designed to detect secreted inhibitory factors showed no interaction between the secondary consumer and primary degraders. Moreover, co-cultures of vsple1A01 and palte3D05 with marib6B07 in liquid media supplemented with #-acetylglucosamine (the monomer of chitin), as sole carbon source showed no decrease in growth rates (Fig S8). This suggests that either an antagonistic factor is only secreted in the particle environment, or what is more likely, that the observed inhibition of primary degrader growth is based on interference with physical processes that only take place when resources are concentrated on particles (e.g. colonization, attachment, etc.).

## Discussion

Despite the significant efforts put into understanding the factors that drive the turnover of organic matter in the ocean^26,27^, the potential role that microbial interactions may play in this process has remained relatively unexplored. Our study leveraged a simplified model based on wild isolates that naturally colonize chitin particles to dissect this question. We provided evidence that both differences in primary degrader type and the ratio of primary degrader to secondary consumer can significantly alter particle degradation kinetics. Remarkably, we show that even in the ideal conditions of our experiments (no N limitation, high number of cells pre-grown in rich media) particle turnover times can be as high as 200 hours or more, that is, in the same range as the residence time of particles in the ocean’s mixed layer. Moreover, we showed interactions between primary degraders and secondary consumers lead to a significant increase in particle turnover times. This result is in agreement with our previous observation of colonization dynamics in natural seawater, which showed that secondary consumers “displace” primary degraders from particles, becoming the dominant members of the particle attached community after a brief initial period of colonization by degraders^16^. Taken together these results suggest that the micro-scale community ecology of particle-attached bacteria plays an important role in controlling rates of carbon turnover in the ocean.

Although in this study we do not identify a direct mechanism for the inhibitory effect of secondary consumers on primary degraders, our results suggest that the effect is not dependent on chemical interactions, which tend to be strain specific. Instead, the fact that we were able to observe degradation inhibition with different secondary consumers in a dose-specific manner suggests that a role of physical processes such as occlusion of the particle surface or an alteration of resource gradients around the particle, which are likely to occur regardless of species identity. Furthermore, this notion is consistent with the fact that degradation inhibition was only observed when bacteria grow on particles, and that the consequences of adding secondary consumers to the environment are similar to those of reducing the primary degrader load (and hence their particle colonization rate). Finally, the fact that the secondary consumer load required to induce degradation inhibition is anticorrelated with the hydrolytic power of the degrader reinforces the notion that particle depolymerization and secondary consumer growth are competing processes. Further work should aim at identifying the precise mechanisms that mediate the negative feedback from secondary consumers to degraders, tracking single cell behavior on and around particles as well as the interplay between spatial structure enzymatic activity.

## Materials and Methods

### Bacterial culturing conditions

Bacterial strains used in this study were previously isolated from model chitin particles^16^. Strains were streaked from glycerol stocks onto Marine Broth 2216 (Difco #279110) 1.5 % agar (BD #214010) plates. After 48 h, single colonies were transferred to 2 ml liquid Marine Broth 2216 and incubated at room temperature, shaking at 200 rpm. Saturated liquid cultures were harvested after 48 h by centrifugation for 8 minutes at 3000 rpm (Eppendorf 5415D, Rotor F45–24–11) and washed two times with Tibbles-Rawling minimal media (see supplemental material of ref ^16^ for a detailed recipe). Optical density (OD) 600 nm was determined in 200 μl (50 μl culture, 150 μl minimal media) in a clear 96-well plate (VWR 10062–900) with a spectrophotometer (Tecan Infinite F500). Cell numbers were normalized to the desired initial concentrations using a three-point linear calibration between OD 600 nm and direct cell counts determined with a Guava easyCyte Benchtop Flow Cytometer for each strain.

### Particle degradation experiments

Particle degradation experiments were performed in clear 96 well plates (VWR 10062–900). Each well contained 180 μl Tibbles–Rawling minimal, bacterial cells at defined concentrations prepared as described above, and approximately 100 chitin magnetic beads (New England Biolabs #E8036L). Before being used in the experiments, the chitin magnetic beads storage buffer was removed using a neodymium magnet (McMaster-Carr #5862K38) to retain the beads. Beads were washed twice in Tibbles–Rawling minimal media and size selected using 100µm and 40μm strainers (VWR, #10199–658 and #10199–654, respectively).

For Fig 1B, the colonized particle was stained in the well after 24 by adding Syto9 (Thermo Fisher, S34854), 500 nM final concentration for 1h at room temperature in the dark. Microscopy was performed on an EVOS FL Auto Imaging System (Fisher #AMAFD1000) using a GFP lightcube (Thermo Fisher AMEP4651) and a 20x fluorite, long working distance objective (Fisher #AMEP4682, NA 0.40, WD 3.1 mm) and the softwares’ (revision 31201) Z-stack function. 3D-reconstruction was done using the ImageJ distribution Fiji (ImageJ 1.51N).

### Time lapse imaging

Phase contrast time lapse images were acquired with an EVOS FL Auto Imaging System (Fisher #AMAFD1000) using the EVOS software (revision 31201) and a 20x fluorite, long working distance, phase-contrast objective (Fisher #AMEP4682, NA 0.40, WD 3.1 mm). Images were manually focused for each particle to capture the maximum cross section area (see Fig 1C, upper panel). Time lapses ran a maximum of 240 h, with images acquired every 2 h. To minimize evaporation effects, culturing plates were wrapped in para film during the time-lapse experiments and outer wells filled with 200 μl water.

### Image processing and Volume quantification

Phase contrast images were analyzed using the ImageJ distribution Fiji (ImageJ 1.51N). A polygonal shape was manually drawn around the particle to determine the area of the particles’ cross-section. To convert from cross section area in square pixel (1 pixel = 0.4545 μm) to volume (in μm^3), we assumed a spherical shape of the particles. Volumes were normalized to initial volume at t=0 h to account for variation in particle sizes. In order to estimate the particle half-life, we fitted a sigmoidal function 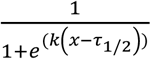 using MATLAB (Version R2016b) and the ‘fit’ function with initial values for *k* (0.5) and *τ*_1/2_ (initial estimates vary for each strain), constraining both variables to positive values (see also Fig S3 for examples of sigmoidal fits to the data).

### Co-culture experiments

Cell counts were obtained by sampling 100 μl from 96 well culture plates (inoculated with 170 μl minimal media, 2x10 μl of the normalized bacterial culture, and 10 μl particles as described above). Imaging was performed as described above. For CFU counts, samples were vortexed thoroughly to detach cells from particles and 10 μl were plated in 10^-2 and 10^-3 dilutions in replicates on MB2216 agar plates using rattler beads (Zymo S1001). After 72 h, colonies were counted to obtain CFUs.

### DNA quantification

To quantify DNA as a proxy for biomass from mono cultures in 96 well plates, wells were mixed thoroughly by pipetting and 100 μl of each well (including the particles) were sampled and frozen at −20 °C for subsequent analysis. Cells were lysed by thawing and boiling (95 °C, 10 min) 10 μl of each sample. Lysed samples were diluted 1:10 in TE buffer and quantified using Quant-it pico green (Fisher # P7589) standard protocols.

### Strain genome annotation

The genomes are deposited at NCBI under Bioproject # PRJNA414740 and the respective accession numbers in Table S1. Assembled genomes were annotated using RAST and genome content was parsed using text parsing of the genome annotations for Chi, ChB, GTx.

## Acknowlegements

The authors wish to thank members of the Cordero lab for thoughtful discussions. This research was supported by NSF grant OCE-1658451, European Starting Grant no. 336938. OXC was also supported by the Simons Early Career Award 410104 and the Alfred P Sloan fellowship FG-20166236.

## Supplementary Material

### Supplementary Text

#### Modeling of particle half-life for non-cooperative degraders

The degradation of a chitin particle by bacteria can be modeled in a simple way by taking into account two processes: (i) Free-living bacteria attach to the particle surface at a rate, *a*, proportional to their planktonic concentration, [B0], such that *a* = *a*_0_[B0], where *a*_0_ is the attachment rate per bacterial cell; (ii) Attached cells degrade the particle at a ratep, and chitin monomers are converted to bacterial biomass at a conversion factor *r*. Note, that the conversion factor *r* may take into account the loss of monomers to the environment. This results in a set of differential equations for the amount of bacterial biomass, B(t), and the total amount of particle, R(t),
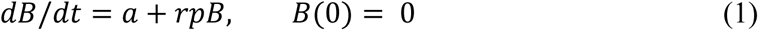

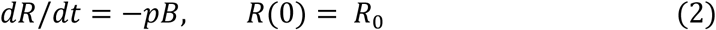

In the above parametrization, the degradation of a chitin particle is described by four independent parameters: (i) the total size of the particle, *R*_0_; (ii) the attachment rate of bacteria, *a*; (iii) the biomass conversion rate, *r*; and (iv) the degradation rate; *p*. There is, however, a more canonical parametrization: Let b(t) = B(t)/*r*, *α* = *a/r* and *β* = *rp*. Then,
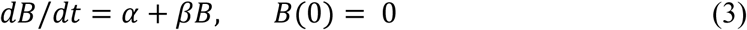

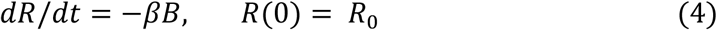

which, for initial conditions *B*(*0*) = 0 and R(0) = *R*_0_, are solved by the equations
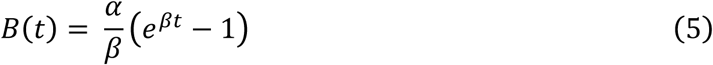

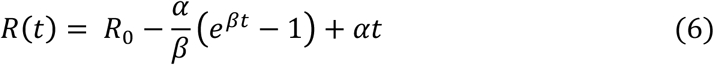

From Eq. 6 the time required to fully degrade the particle can be found by numerically solving the transcendental equation for *R*( *T*) *= R*_0_. Additional analytical insight can be gained, however, by assuming that attachment is slow compared to growth, and hence *R*(*T*) *= R*_0_ *− B*(*T*). Furthermore, we additionally assume that *rpT* ≫ 0 and hence ln(e*^rpT^* − 1) ≈ *rpT*. This leads to a simple expression for the total degradation time, *T*, required to fully degrade a particle,
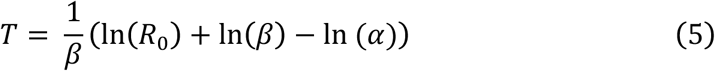

Hence, the degradation time T depends linearly on the logarithm of the attachment rate *a*, and hence the planktonic concentration of bacteria (see Fig. S4)

#### Expected half-life for cooperative degraders

To calculate half-lives in the presence of cooperative growth we used the simple model
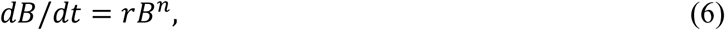

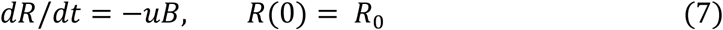

Unfortunately it is not practical to work with the analytical solutions of these set of equations, so we turn to numerical simulations in which we calculate the time it takes to consume half of the resources (*R*(*τ*_1/2_) = *R*(0)/2). For this simulations we use *r =* 0.01 and *u =* 0.5, and explore the shape of the *B*(0) vs *τ*_1/2_ relation by fitting a log linear or a power law relationship. We find that quickly as n > 1 the relationship converges to *τ*_1/2_ ~ 1/*B*(0). Simulations were performed in R using the dSolve package.

## Supplementary Figures

**Figure S1:**
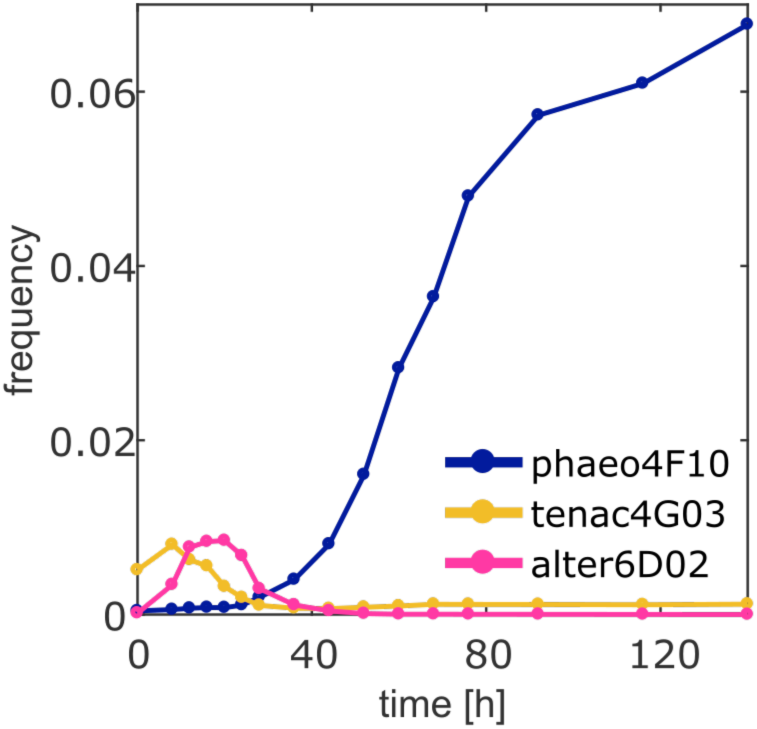
Culture independent dynamics of secondary consumers. phaeo4F10, tenac4G03 and alter6D02. Trajectories shown depict dynamics of selected taxa from particle incubations with raw seawater, where other taxa were present. Data from ^16^.

**Figure S2:**
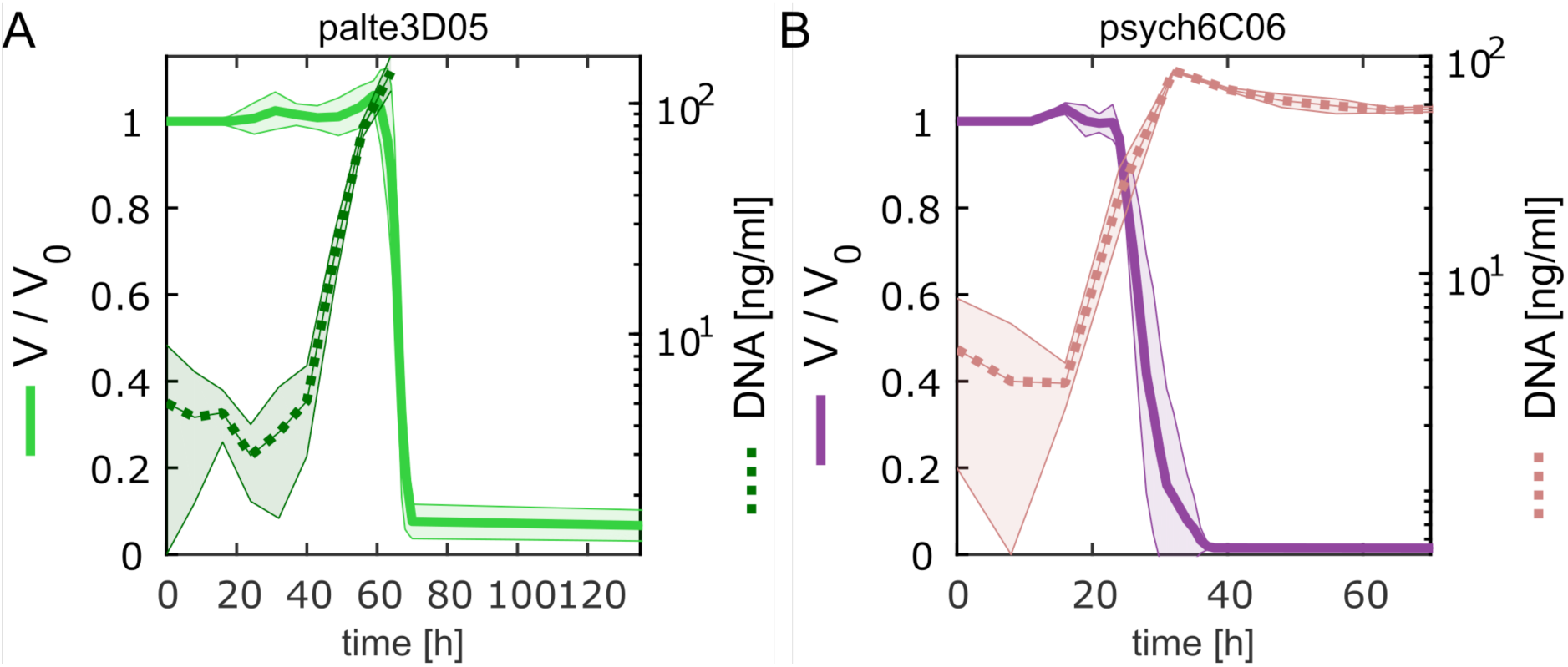
Degradation dynamics and bacterial growth for palte3D05 and psych6C06. Particle volume over time normalized to initial volume (solid line) and bacterial abundance as measured by the amount of DNA extracted from ~100 particles at different points of colonization (dashed line). The standard deviation of measurements was calculated using three replicate particles from the same well, and three different bulk incubations for DNA.

**Figure S3:**
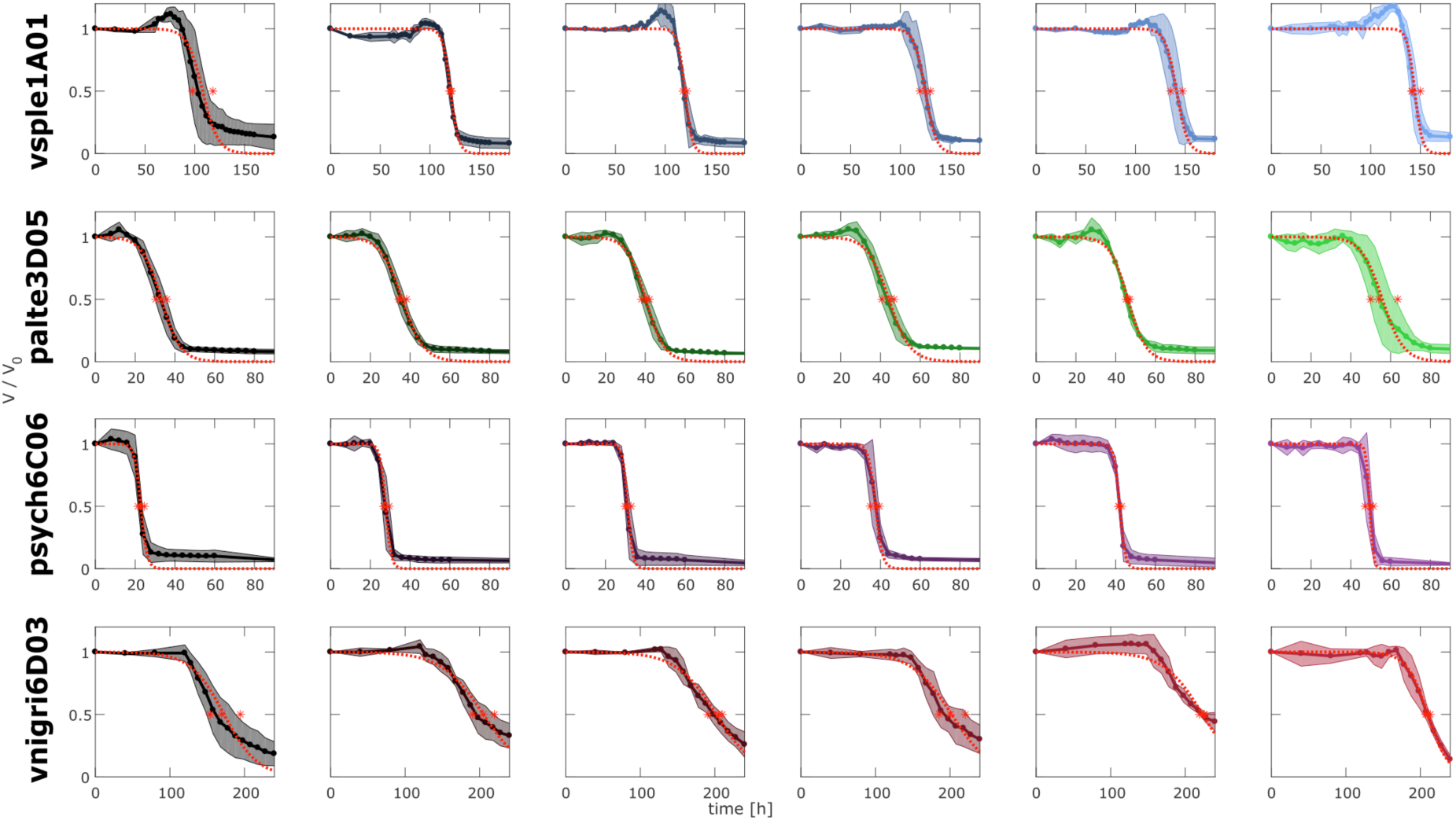
Particle volume over time for different initial concentration of primary degraders. Data corresponds to Fig 2 A-D. Shown are quantified, normalized particle volumes for four primary degraders and six initial cell concentrations (from left to right: 2^10, 2^8, 2^6, 2^4, 2^2, 2^0 cells / μl). Solid line: mean, shaded area: standard deviation of n=3 replicate particles. Dashed red line: fit of a sigmoidal function to the mean; red asterisk: inferred tau for the three single replicates (see methods).

**Figure S4:**
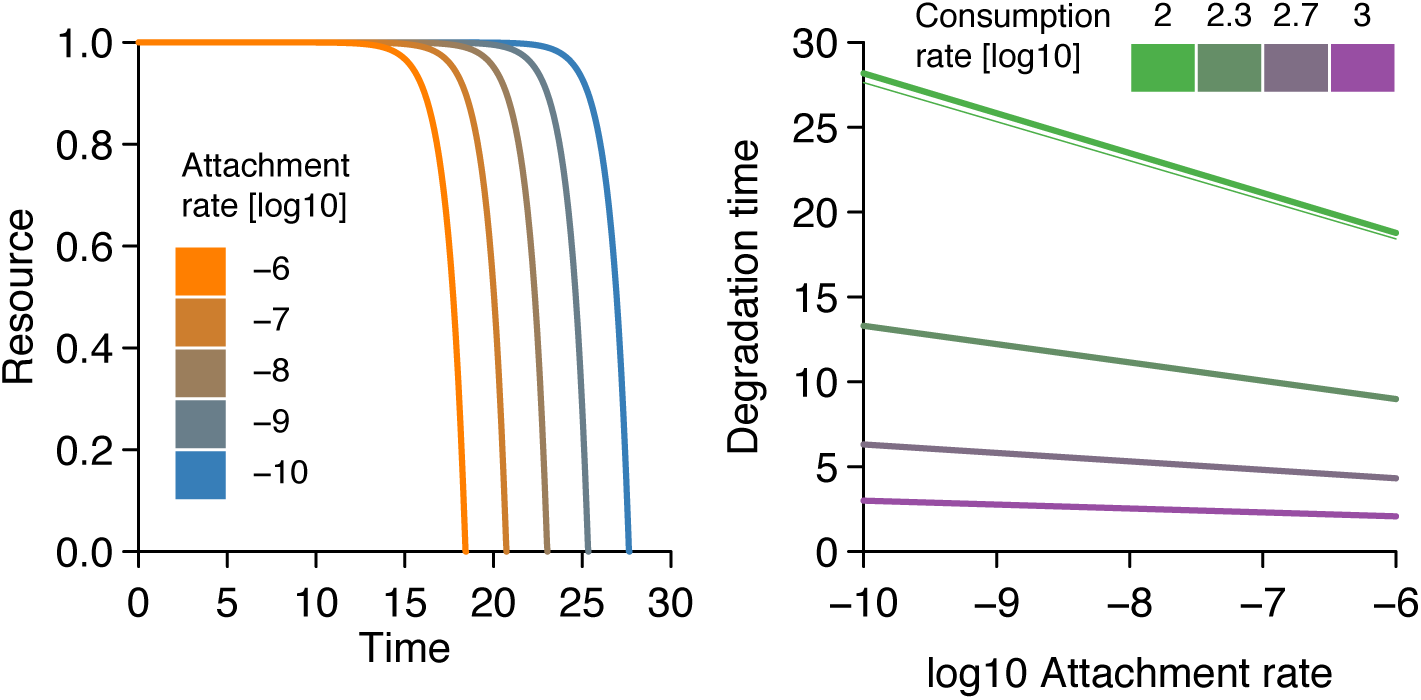
Left panel: degradation dynamics predicted by equations 1–6. Attachment rate is the product of the per-cell attachment rate and the number of initial bacterial in the medium. Right panel: particle half-lives as a function of attachment rates for populations with different hydrolytic powers

**Figure S5:**
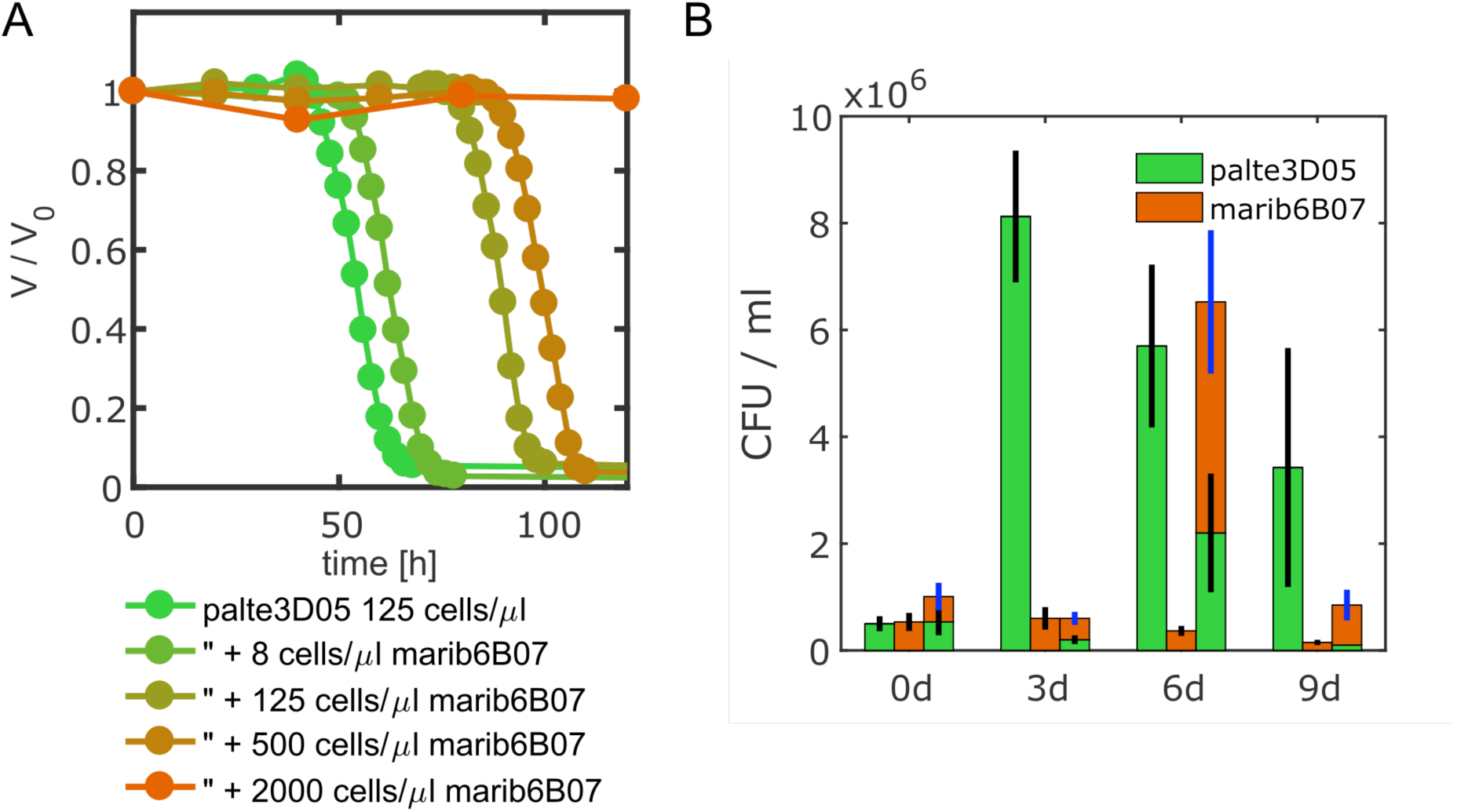
Secondary consumer can inhibit degradation of primary degrader palte3D05. **A**) Particle degradation curves with different marib6b07 concentrations. At increasing concentrations of the secondary consumer the particle half-life increases disproportionally beyond the 220 h time limit (endpoint not shown). **B**) CFUs of palte3D05 (500 cells / μl) and marib6B07 (500 cells / μl) during mono- (first two bars for each time point) and co-culture (third, stacked bar) on chitin particles, showing that palte3D05 growth in mono-culture peaks at ~3d and marib6B07 cannot grow in mono-culture. Marib6B07 grows “at the expense” of palte3D05’s yield and delays peak growth and particle degradation which occurred at ~6d. Black error bars correspond to palte3D05, blue error bars to marib6B07, respectively (both depict standard deviation for n=3 replicates). Decrease in CFUs is due to loss of viability after degradation.

**Figure S6:**
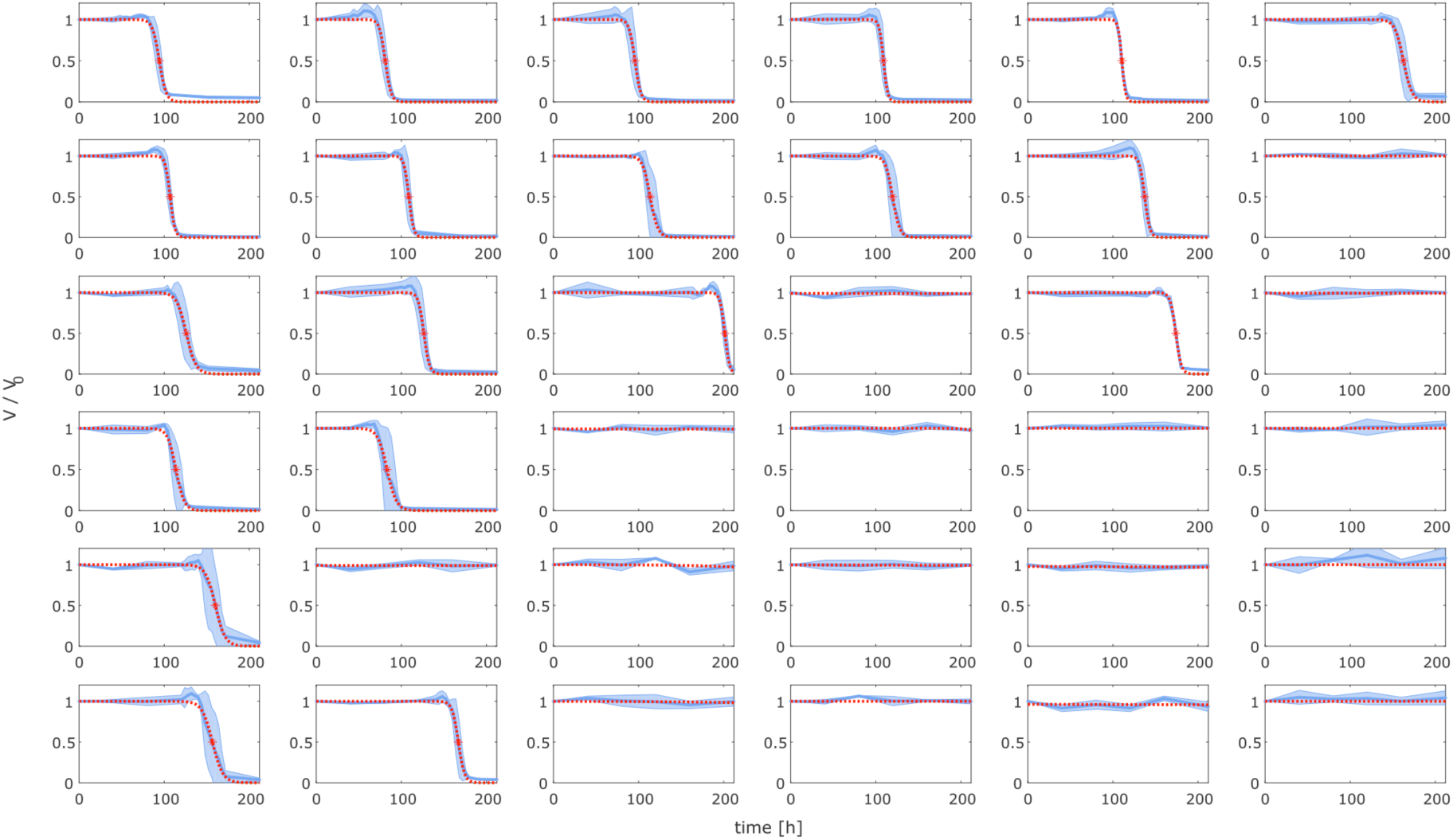
Particle volume over time for different initial concentrations of primary degrader vsple1A01 and secondary consumer marib6B07. Data corresponds to Fig 3C, left, heatmap. Shown are quantified, normalized particle volumes for all fields of the heat map in the same arrangement. Solid line: mean, shaded area: standard deviation of n=3 replicate particles. Dashed red line: fit of a sigmoidal function to the mean; red asterisk: inferred tau from mean as shown in heat map (see methods).

**Figure S7:**
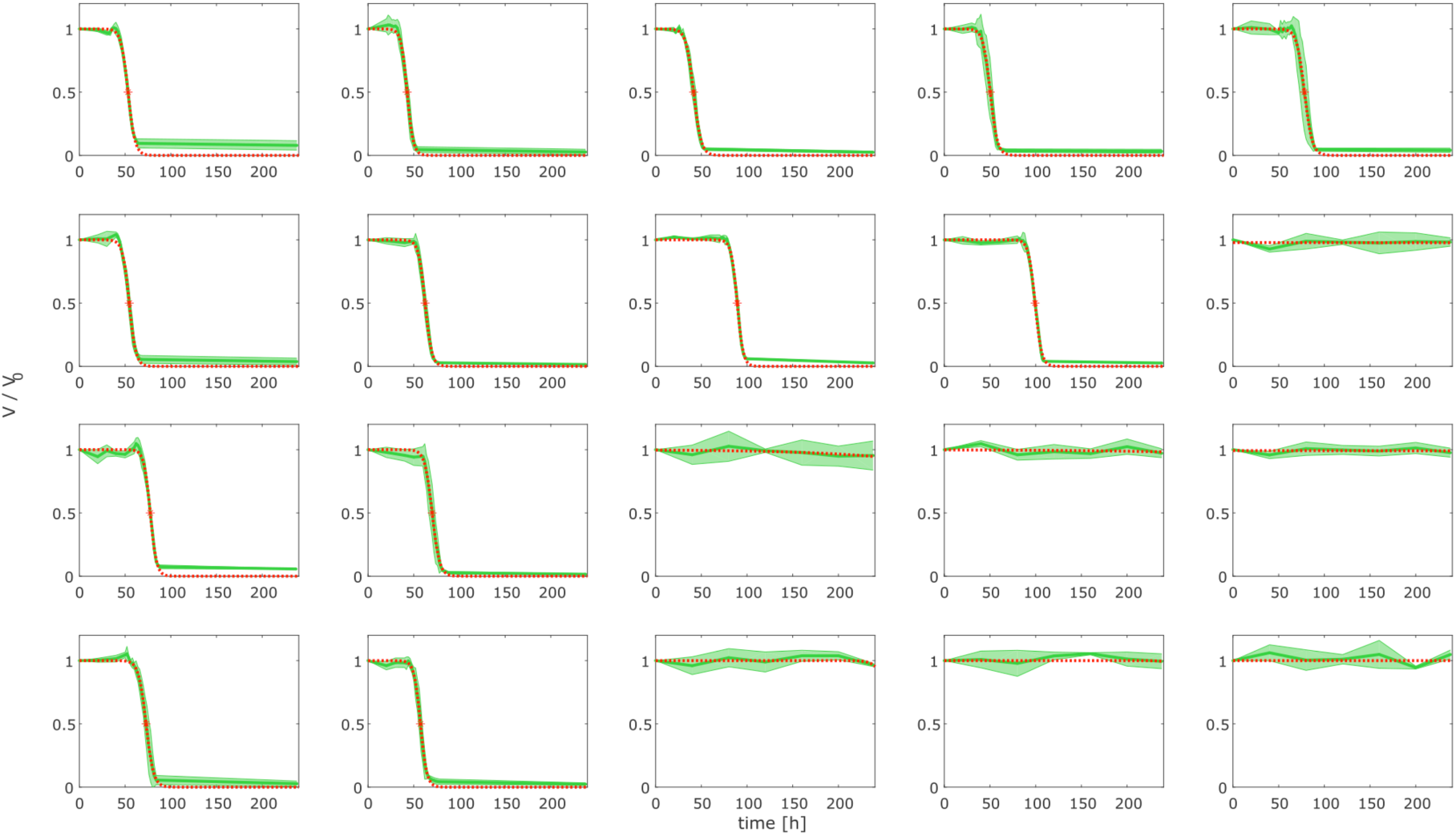
Particle volume over time for different initial concentrations of primary degrader palte3D05 and secondary consumer marib6B07. Data corresponds to Fig 3C, right heatmap. Shown are quantified, normalized particle volumes for all fields of the heat map in the same arrangement. Solid line: mean, shaded area: standard deviation of n=3 replicate particles. Dashed red line: fit of a sigmoidal function to the mean; red asterisk: inferred tau from mean as shown in heat map (see methods).

**Figure S8.**
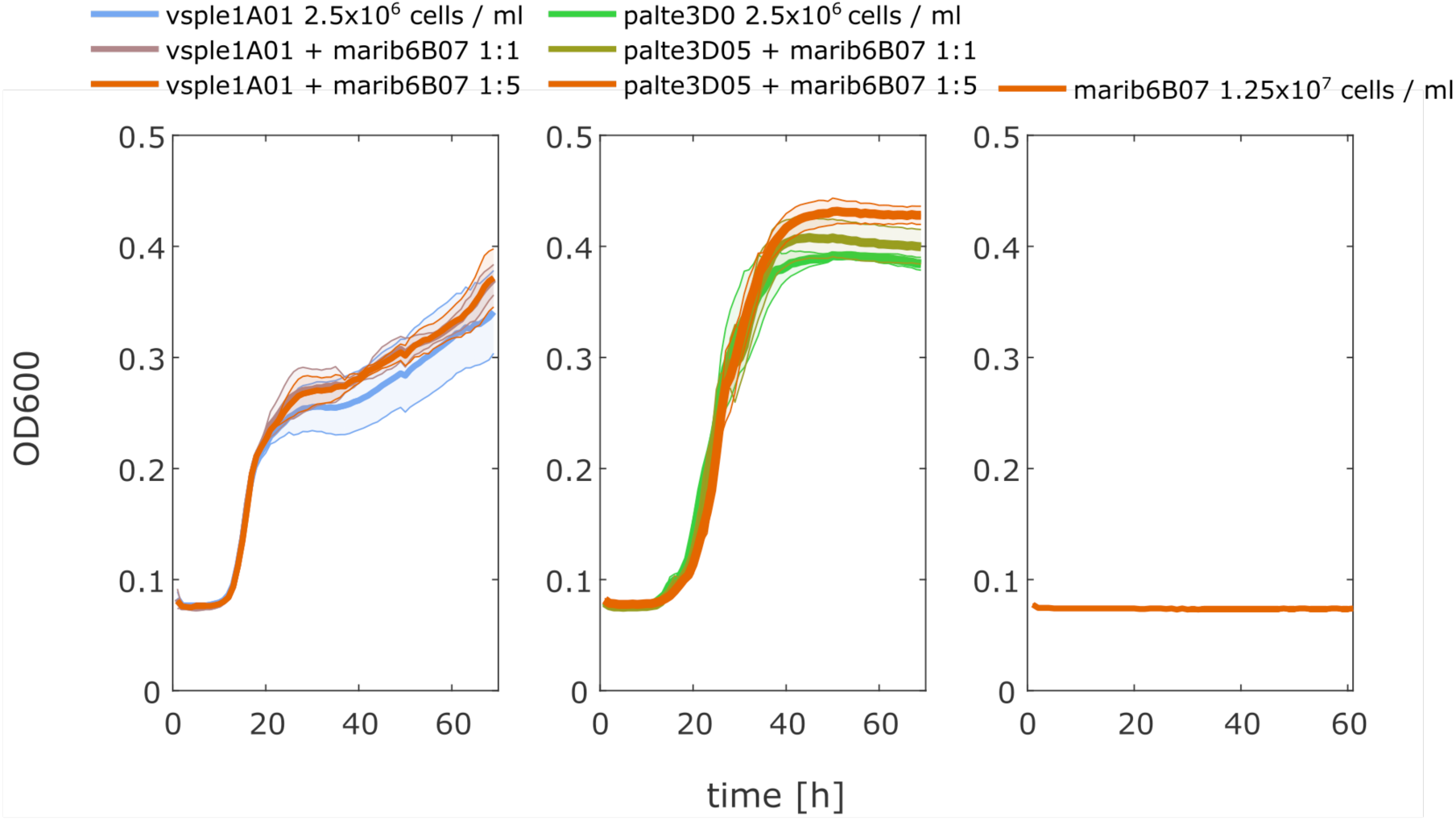
Growth of secondary consumer marib6B07 in co-culture with primary degrader vsple1A01. (left panel), palte3D05 (middle), and in monoculture (right panel) on 0.1 % GlcNAc (A-Acetyglucosamin, chitin monomers). Co-cultures are in 1:1 and 1:5 ratios of primary degrader to secondary consumer, as indicated above the panels.

**Figure S9:**
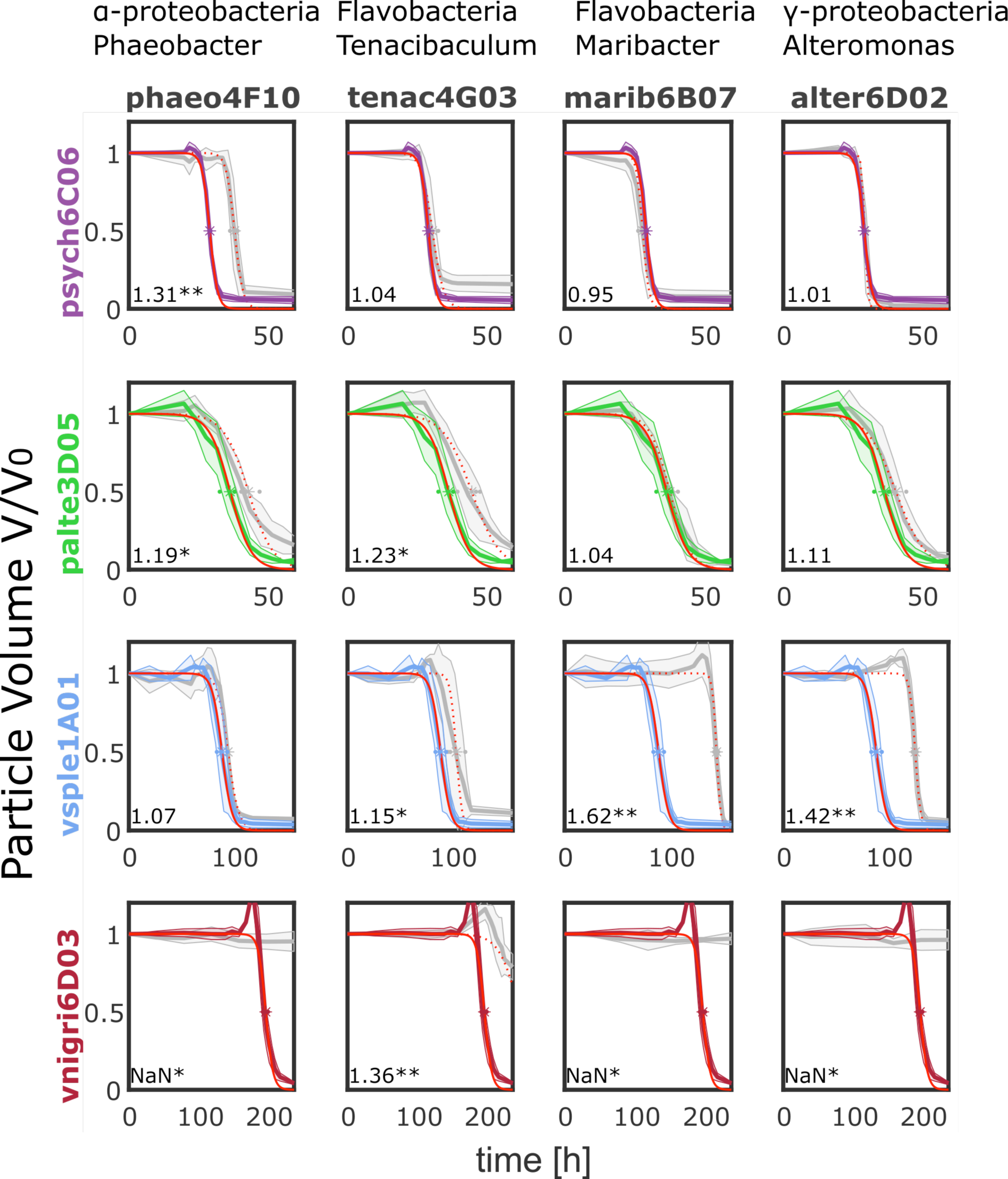
Data underlying the network in Fig. 4. Each panel depicts the normalised particle degradation over time for the primary degrader (row) and the secondary degrader (colum) in the respective colors indicated on the strain name. Solid line: mean, shading: standard deviation, n=3. Red solid line: sigmoidal fit to infer *τ*_1/2_ for the respective primary degrader, red dashed line sigmoidal fit to infer *τ*_1/2_ for the respective co-culture. Asterisk indicates the values of each replicate for *τ*_1/2_. Text bottom left indicates the ratio of to 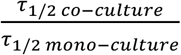 used to infer the edge thickness of the network in figure 4 and black asterisks indicate significance levels, * < 0.05, ** < 0.1. See also Table S3.

**Supplemental Movie 1. 3D reconstruction of a chitin micro particle** colonized by palte3D05 for 24 h, stained with SYTO9.

**Supplemental Movie 1. Phase contrast time-lapse** of a chitin particle cross section taken during degradation by vsple1A01, corresponding to frames shown in Figure 1C.

## Supplementary Tables

**Table S1:**
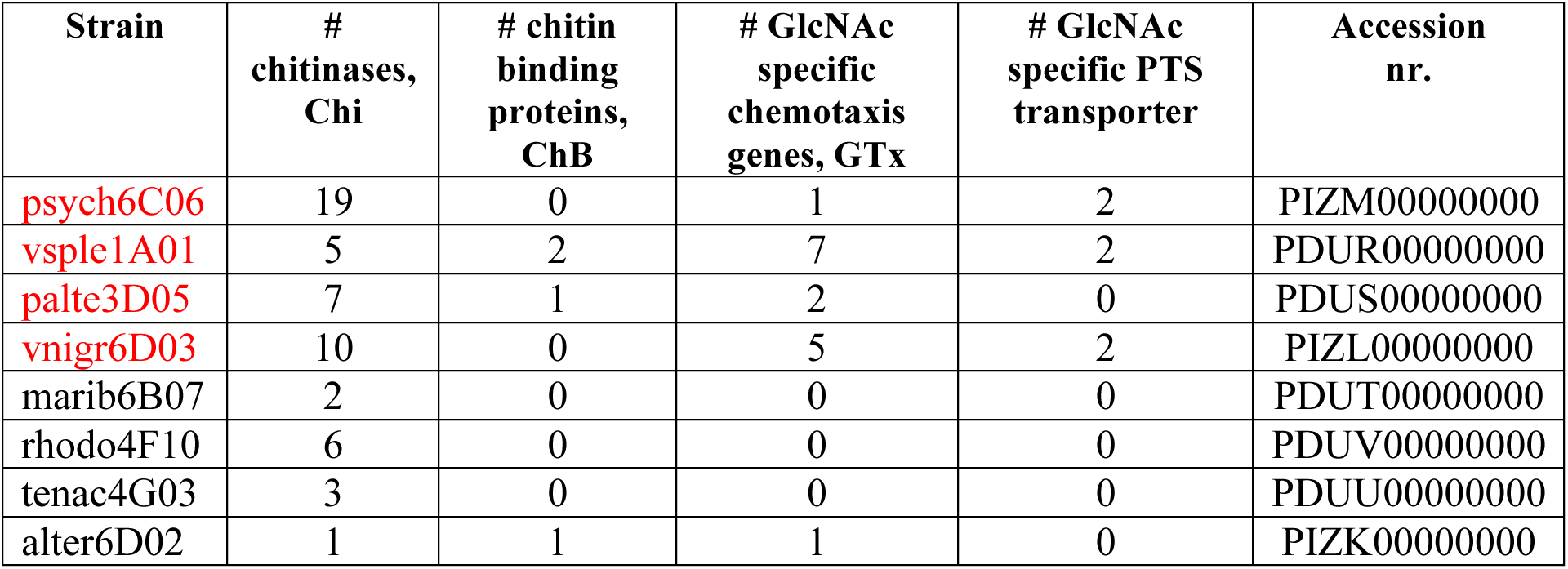
Genomic features of chitin degraders (in red) and non-degraders. Chitin degraders tend to have multiple copies of chitinases, as well as chitin binding proteins, GlcNAc chemotaxis and PTS transporter genes. The genomes are deposited at NCBI under Bioproject # PRJNA414740 and the respective accession numbers below.

**Table S2:** R^2^ and p-value of the multiple linear regression from Figure 2E.

| | $R^2$ | p-value |
| --- | --- | --- |
| psych6C06 | 0.96 | 5.63E-13 |
| palte3D05 | 0.82 | 1.97E-07 |
| vsple1A01 | 0.78 | 1.46E-05 |
| vnigri6D03 | 0.40 | 4.88E-03 |

## References

1. Deleersnijder, E., Beckers, J.-M. & Delhez, E. J. M. The Residence Time of Settling Particles in the Surface Mixed Layer. Environ. FluidMech. 6, 25–42 (2006).

2. Loreau, M. Biodiversity and Ecosystem Functioning: Current Knowledge and Future Challenges. Science (80-.). 294, 804–808 (2001).

3. Balvanera, P. et al. Quantifying the evidence for biodiversity effects on ecosystem functioning and services. Ecol. Lett. 9, 1146–1156 (2006).

4. Hector, A. & Bagchi, R. Biodiversity and ecosystem multifunctionality. Nature 448, 188–190 (2007).

5. Cordero, O. X. & Datta, M. S. Microbial interactions and community assembly at microscales. Curr. Opin. Microbiol. 31, 227–234 (2016).

6. Volkman, J. K. & Tanoue, E. Chemical and Biological Studies of Particulate Organic Matter in the Ocean. J. Oceanogr. 58, 265–279 (2002).

7. Passow, U. Transparent exopolymer particles (TEP) in aquatic environments. Prog. Oceanogr. 55, 287–333 (2002).

8. Alldredge, A. L. & Silver, M. W. Characteristics, dynamics and significance of marine snow. Prog. Oceanogr. 20, 41–82 (1988).

9. Engel, A., Thoms, S., Riebesell, U., Rochelle-Newall, E. & Zondervan, I. Polysaccharide aggregation as a potential sink of marine dissolved organic carbon. Nature 428, 929–32 (2004).

10. Andrew McDonnell and Ken Buesseler. Marine Particle Dynamics: Sinking Velocities, Size Distributions, Fluxes, and Microbial Degradation Rates: Recent Dissertations and Theses: MIT/WHOI Joint Program. PhD Thesis (2011).

11. Yu, C., Lee, A. M., Bassler, B. L. & Roseman, S. Chitin utilization by marine bacteria. A physiological function for bacterial adhesion to immobilized carbohydrates. J. Biol. Chem. 266, 24260–7 (1991).

12. Stocker, R., Seymour, J. R., Samadani, A., Hunt, D. E. & Polz, M. F. Rapid chemotactic response enables marine bacteria to exploit ephemeral microscale nutrient patches. Proc. Natl. Acad. Sci. 105, 4209–4214 (2008).

13. Stocker, R. Marine microbes see a sea of gradients. Science 338, 628–33 (2012).

14. Long, R. A. et al. Antagonistic interactions among marine bacteria impede the proliferation of Vibrio cholerae. Appl. Environ. Microbiol. 71, 8531–6 (2005).

15. Long, R. A. & Azam, F. Antagonistic interactions among marine pelagic bacteria. Appl. Environ. Microbiol. 67, 4975–4983 (2001).

16. Datta, M. S., Sliwerska, E., Gore, J., Polz, M. F. & Cordero, O. X. Microbial interactions lead to rapid micro-scale successions on model marine particles. Nat. Commun. 7, 11965 (2016).

17. Jagmann, N., von Rekowski, K. S. & Philipp, B. Interactions of bacteria with different mechanisms for chitin degradation result in the formation of a mixed-species biofilm. FEMSMicrobiol. Lett. 326, 69–75 (2012).

18. Fontanez, K. M., Eppley, J. M., Samo, T. J., Karl, D. M. & DeLong, E. F. Microbial community structure and function on sinking particles in the North Pacific Subtropical Gyre. Front. Microbiol. 6, 469 (2015).

19. Whitman, W. B., Coleman, D. C. & Wiebe, W. J. Prokaryotes: The unseen majority. Proc. Natl. Acad. Sci. 95, 6578–6583 (1998).

20. T. K. L. Meyvis, S. C. De Smedt, * and, Demeester, J. & Hennink, W. E. Influence of the Degradation Mechanism of Hydrogels on Their Elastic and Swelling Properties during Degradation. (2000). doi:10.1021/MA992131U

21. Drescher, K., Nadell, C. D., Stone, H. A., Wingreen, N. S. & Bassler, B. L. Solutions to the public goods dilemma in bacterial biofilms. Curr. Biol. 24, 50–5 (2014).

22. Kovárová-Kovar, K. & Egli, T. Growth kinetics of suspended microbial cells: from single-substrate-controlled growth to mixed-substrate kinetics. Microbiol. Mol. Biol. Rev. 62, 646–66 (1998).

23. Le Roux, F. et al. Genome sequence of *Vibrio splendidus*: an abundant planctonic marine species with a large genotypic diversity. Environ. Microbiol. 11, 1959–1970 (2009).

24. Lara, E. et al. Life-Style and Genome Structure of Marine Pseudoalteromonas Siphovirus B8b Isolated from the Northwestern Mediterranean Sea. PLoS One 10, e0114829 (2015).

25. Romanenko, L. A. et al. Pseudoalteromonas agarivorans sp. nov., a novel marine agarolytic bacterium. Int. J. Syst. Evol. Microbiol. 53, 125–131 (2003).

26. Jiao, N. et al. Microbial production of recalcitrant dissolved organic matter: long-term carbon storage in the global ocean. Nat. Rev. Microbiol. 8, 593–9 (2010).

27. Kirchman, D. L. Microbial Ecology of the Oceans, 2nd Edition – David L. Kirchman.

